# Adeno-associated viruses for efficient gene expression in the axolotl nervous system

**DOI:** 10.1101/2024.02.15.580426

**Authors:** Katharina Lust, Elly M. Tanaka

## Abstract

Axolotls are models for studying nervous system evolution, development, and regeneration. Tools to visualize and manipulate cells of the axolotl nervous system with high efficiency, spatial and temporal precision are therefore greatly required. Recombinant adeno-associated viruses (AAVs) are frequently used for *in vivo* gene transfer of the nervous system but virus-mediated gene delivery to the axolotl nervous system has not yet been described. Here, we demonstrate the use of AAVs for efficient gene transfer within the axolotl brain and the retina. We show that serotypes AAV8, AAV9, AAVRG and AAVPHP.eB are suitable viral vectors to infect both excitatory and inhibitory neuronal populations of the axolotl brain. We further use AAV9 to trace retrograde and anterograde projections between the retina and the brain and identify a cell population projecting from the brain to the retina. Together, our work establishes AAVs as a powerful tool to interrogate neuronal organization in the axolotl.

## Introduction

Axolotls (*Ambystoma mexicanum*) can regenerate injuries to the central and peripheral nervous system making them an ideal model to study formation and regeneration of neuronal circuits. The delineation of neuronal circuits in salamanders has primarily relied on classical tracing techniques, employing for instance Golgi staining, horse radish peroxidase, Neurobiotin, and biotinylated dextran amine^1–4^. Tools that will allow for efficient and functional dissection of neuronal circuitry are still lacking in salamanders.

While transgenesis^5^, electroporation^6^ and virus-mediated gene delivery^7^ are established methodologies for introducing genes into salamander cells. Several virus-mediated gene delivery tools including pseudotyped Moloney murine leukemia virus^8^, vaccinia virus^9^, foamy virus^10^, and pseudotyped baculovirus^7^ were shown to be functional and efficient for transduction of cells in the axolotl limb where they infect muscle, cartilage, dermis, fibroblasts and Schwann cells^7,10^. Virus-mediated gene delivery to axolotl nervous system tissue has not been achieved so far. Neuron-targeted gene delivery in the central and peripheral nervous system is challenging because neurons are postmitotic. For that purpose, Adeno-associated viruses (AAVs) and lentiviruses, are commonly used viral vectors in neuroscience^11,12^. While lentivirus has been utilized in axolotls, the observed expression levels are notably low. Especially *in vivo* hardly any expression of lentivirus-mediated transgenesis has been observed^10^. Use of AAVs has so far not been described in the axolotl.

AAVs are particularly attractive candidate vectors because in contrast to most viruses they possess no intrinsic pathogenicity, low immunogenicity, long transgene expression and a wide-ranging tropism^11^. The utilization of AAV-mediated delivery for introducing genetic material into specific populations of the nervous system represents a valuable strategy for investigating both anatomy and circuit function^11^. By entering either at somata or axon terminals, anterograde and retrograde projections can be assessed allowing to map input and output projections of defined regions^13^. AAV vectors, widely employed in rodents and other mammals^14^, have recently found application in avian species such as pigeon^15^ and songbirds^16,17^, as well as in the reptile *Pogona vitticeps*^18^. In contrast, AAVs fail to infect neurons in zebrafish^19^ and show only limited efficiency in *Xenopus laevis*^20^. Several studies have conducted comparative evaluations of AAV serotypes and it has been demonstrated that the tropism of AAVs for distinct organs and tissues varies depending on their serotype^21^.

Importantly, the transduction efficacy of a serotype is not simply predictable from one model system to another.

Here, we present a set of AAV serotypes with high transfection efficiency for the axolotl brain and retina. In short, we tested seven AAV serotypes for their infection and expression efficiency and found that AAV8, AAV9, AAVRG and AAVPHP.eB are most suited to label neurons of the brain. We also investigate the use of AAVs to label input and output projections of the retina through anterograde and retrograde tracing. We define AAV9 as the most suitable AAV to label retinal cells and use it to to trace projections from the retina to various brain regions. Additionally, we identify a retinopetal cell population in the axolotl which projects from the brain to the retina that had not been documented in this salamander species before.

## Results

### Comparative transduction analysis of AAV serotypes in the axolotl brain

AAV serotypes have been determined to infect different organs in mammalian model systems with various efficiencies^21^. As the use of AAVs have not yet been described in the axolotl we first explored which serotypes can efficiently infect cells of their central nervous system. For this we tested seven commonly used AAV serotypes: AAV1, AAV2, AAV5, AAV8, AAV9, AAVRG and PHP.eB. To distinguish lack of transduction from lack of expression we used the same transgene CAG:GFP in each virus with a titer ≥ 7×10¹² vg/mL. We injected the optic tectum of ten animals of a size of 3 cm (nose to tail) with each serotype and observed them under a fluorescent widefield microscope for a time period of four weeks (Figure 1A, Figure S1). Two weeks after injection the first brains started to show weak green fluorescence (data not shown) which increased until four weeks post injection (Figure S1). Notably, AAV8-CAG, AAV9-CAG and AAVPHP.eB-CAG displayed strong expression in the brain when observed in live animals (Figure S1). In contrast, expression of GFP was barely detectable in live animals transfected with AAV2-CAG (one out of ten animals positive, Figure 1, Figure S1C). At four weeks post-injection we harvested the brains and performed wholemount immunostaining for GFP. We aimed to determine whether brains with low expression were devoid of GFP-expressing neurons (i.e. lack of infection) or whether the expression of GFP was low and could be amplified with immunostaining. We found abundant GFP-expressing neurons in the optic tectum transduced with each serotype except AAV2 (six neurons in one out of ten brains). We then quantified the density of GFP-positive neurons in a defined volume of 200×200×80µm^3^ and found that AAV8-CAG, AAV9-CAG and AAVPHP.eB-CAG showed the highest efficiency to label cells around the injection site. Especially AAVPHP.eB-CAG showed a high density of labeled cells which reached almost twice as much as AAV8-CAG and AAV9-CAG. We have previously used axolotls of a larger size (10 cm nose to tail) for studying brain regeneration^3^. We therefore also tested the feasibility and efficiency of AAV-mediated transduction in these animals. We decided to use only the AAVs which had worked best in small axolotls and therefore injected AAV8-CAG:GFP, AAV9-CAG:tdTomato, AAVRG-CAG:tdTomato as well as AAVPHP.eB-CAG:GFP into one hemisphere of the telencephalon of three animals each (Figure S2). Four weeks after injection, we fixed the brains and analyzed by wholemount immunostaining for transgene expression. We found that all four tested serotypes infected cells in 10 cm axolotl brains (Figure S2E). However, the density of expressing cells was lower than in 3 cm animals and more variable across different animals (Figure S2F) which could be due to a different neuronal density in bigger animals.

**Figure 1.**
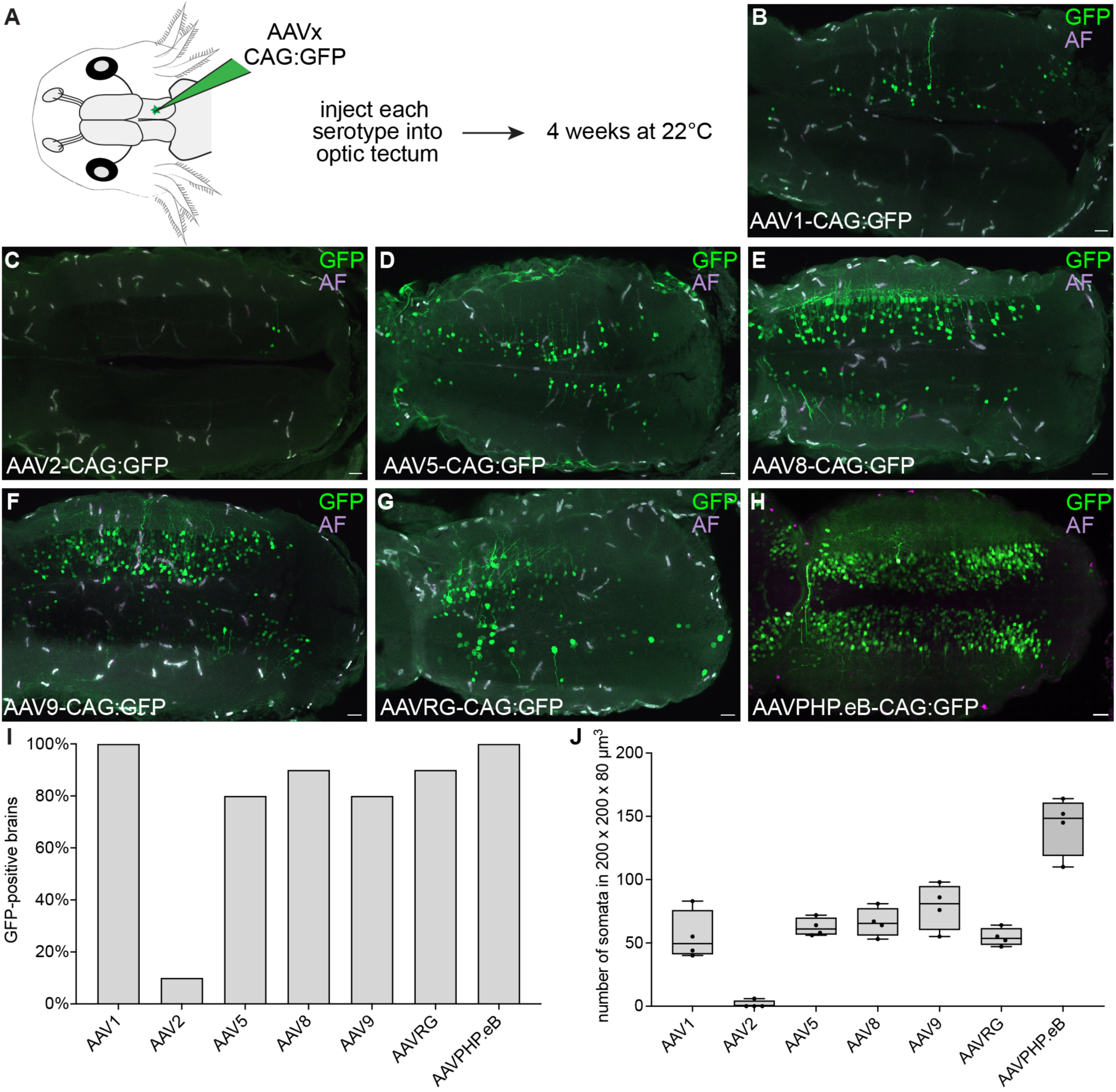
AAV-mediated cell transduction in the larval axolotl brain. (A) Schematic of experimental paradigm for AAV serotype efficiency test. Each serotype was injected into the optic tectum neuropil and animals were housed at 22 degrees for 4 weeks until the analysis. (B-H) Immunofluorescence for GFP (green) and autofluorescence (AF, magenta-white) on wholemount, cleared brains at 4 weeks after AAV injection. All tested AAV serotypes but AAV2 efficiently transduce neurons. Scale bars are 100µm. (n = 10 axolotl for each serotype). (I) Quantification of GFP-positive brains 4 weeks after each AAV serotype injection shows that all tested AAV serotypes but AAV2 efficiently transduce axolotl brains. (n = 10 axolotl for each serotype) (J) Quantification of GFP-positive somata per 200×200×80 µm^3^ shows that AAV8, 9 and PHP.eB transduce the highest number of cells per area. Boxplot indicates “minimum,” first quartile, median, third quartile and “maximum”. (n = 4 axolotl for each serotype)

Taken together, all tested AAV serotypes except AAV2 can be used to deliver genes into brains of larval and juvenile axolotls. AAV8-CAG, AAV9-CAG and AAVPHP.eB-CAG showed the highest efficiency to label cells around the injection site.

### Cellular tropism of AAV8, AAV9 and PHP.eB

The two most abundant cell types in the axolotl brain are neurons and ependymoglia^3^. The CAG promoter can drive transgene expression in all cell types and has been used previously in axolotl for ubiquitous transgene expression^22^. We therefore analyzed the extent to which GFP-expressing cells were co-localized with either a glial or a neuronal marker in sections of brains infected with AAV8-CAG:GFP, AAV9-CAG:GFP and AAVPHP.eB-CAG:GFP. Therefore, we performed combined immunohistochemical stainings on cryosections against GFP to visualize transduced cells, against glial fibrillary acidic protein (GFAP) to visualize ependymoglia and against HuC/HuD to visualize neurons. GFAP and HuC/D staining accounts for the large majority of cells and therefore allow to determine AAV serotype tropism We found that AAV8-CAG (Figure 2A-B), AAV9-CAG (Figure 2C-D) and AAVPHP.eB-CAG (Figure 2E-F) showed significantly higher transgene expression in neurons than in ependymoglia cells, with a maximum of four ependymoglia cells labeled per 18µm section (Figure 2A-F). Thus, even though not specifically targeted to express in neurons AAV8-CAG, AAV9-CAG and AAVPHP.eB-CAG show a higher tropism for neuronal expression. Next, we also investigated the ability of AAV8-CAG, AAV9-CAG and AAVPHP.eB-CAG to transduce excitatory versus inhibitory neurons of the axolotl brain (Figure 2G-I). To facilitate identification of excitatory and inhibitory neurons we used hybridization chain reaction (HCR) in situ for Solute carrier family 17 member 7 *(Slc17a7)* and *Glutamate decarboxylase 2 (Gad2).* We found both GFP, *Slc17a7-*double positive and GFP, *Gad2-*double positive neurons in each infection (Figure 2G-I), indicating that AAV8-CAG, AAV9-CAG and AAVPHP.eB-CAG can infect and drive transgene expression in both excitatory and inhibitory neurons in the axolotl brain.

**Figure 2.**
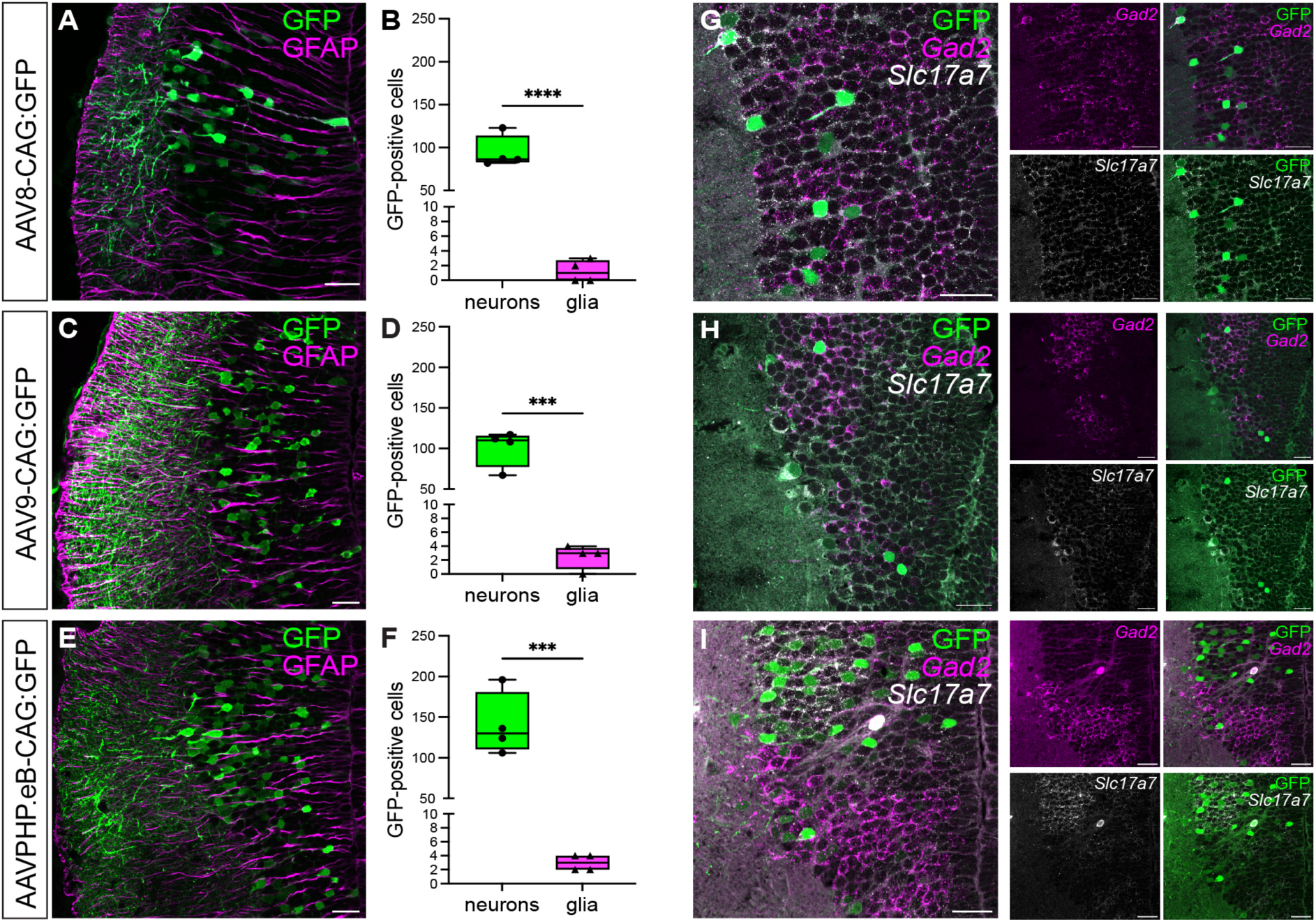
Cellular tropism of AAV8-CAG, AAV9-CAG and AAVPHP.eB-CAG. (A) Confocal image of expression of AAV8-CAG:GFP in neurons versus ependymoglia revealed by immunostaining against GFP (green) and GFAP (magenta). (B) Quantification of GFP positive neurons versus glia in AAV8-CAG:GFP transduced brains. Boxplot indicates “minimum,” first quartile, median, third quartile and “maximum”. Statistical significance is calculated using unpaired t test, ^∗∗∗∗^p < 0.0001. (n = 2 axolotl, quantification of 2 sections per animal) (C) Confocal image of expression of AAV9-CAG:GFP in neurons versus ependymoglia revealed by immunostaining against GFP (green) and GFAP (magenta). (D) Quantification of GFP positive neurons versus glia in AAV9-CAG:GFP transduced brains. Boxplot indicates “minimum,” first quartile, median, third quartile and “maximum”. Statistical significance is calculated using unpaired t test, ^∗∗∗^p <0.0001. (n = 2 axolotl, quantification of 2 sections per animal) (E) Confocal image of expression of AAVPHP.eB-CAG:GFP in neurons versus ependymoglia revealed by immunostaining against GFP (green) and GFAP (magenta). (F) Quantification of GFP positive neurons versus glia in AAVPHP.eB-CAG:GFP transduced brains. Boxplot indicates “minimum,” first quartile, median, third quartile and “maximum”. Statistical significance is calculated using unpaired t test, ^∗∗∗^p < 0.004. (n = 2 axolotl, quantification of 2 sections per animal) (G-I) Confocal image of expression of AAV8-CAG:GFP, AAV9-CAG:GFP and AAVPHP.eB-CAG:GFP respectively in excitatory neurons versus inhibitory neurons revealed by HCR against *Gad2* (magenta) and *Slc17a7* (white). All scale bars are 50 µm.

### Immune cell infiltration and glial activation after AAVPHP.eB-mediated neuronal transduction

Inflammation and glial cell activation associated with AAV administration have been observed after AAV delivery to the brain and retina and depend on many factors, including the AAV serotype, AAV dose, and AAV quality and nature of the transgene^23,24^. To investigate whether inflammation and glial activation occurs after transduction of axolotl neurons we infected brains with either AAV8-CAG-GFP, AAV9-CAG:GFP, AAVRG-CAG:GFP and AAVPHP.eB-CAG:GFP and waited for four weeks to reach high expression. We then performed stainings against GFP, Ionized calcium binding adaptor molecule 1 (Iba1) and glial fibrillary acidic protein (GFAP) on cryosections (Figure 3). Iba1 is a microglia/macrophage-specific calcium-binding protein which can be used to determine their activation and infiltration^25^ while GFAP, a marker for ependymoglia cells in the axolotl brain^3^, is used to determine astrocyte activation after stress or injury to the brain^26^. We found that brains infected with AAV8-CAG:GFP (Figure 3A), AAV9-CAG:GFP (Figure 3B) and AAVRG-CAG:GFP (Figure 3C) only display low numbers of Iba1-positive cells (on average 4 to 9 Iba1-positive cells per 18 µm section). In contrast to that, brains transduced with AAVPHP.eB-CAG:GFP showed strongly increased infiltration of Iba1-positive cells when compared to all other brains (on average 38 Iba1-positive cells per 18 µm section, Figure 3D). However, the expression of GFAP was not significantly changed between all injected serotypes (Figure S3) indicating that ependymoglia cell activation did not occur after AAV transduction. Together, this shows that only AAVPHP.eB-CAG induces a strong increase of immune cells.

**Figure 3:**
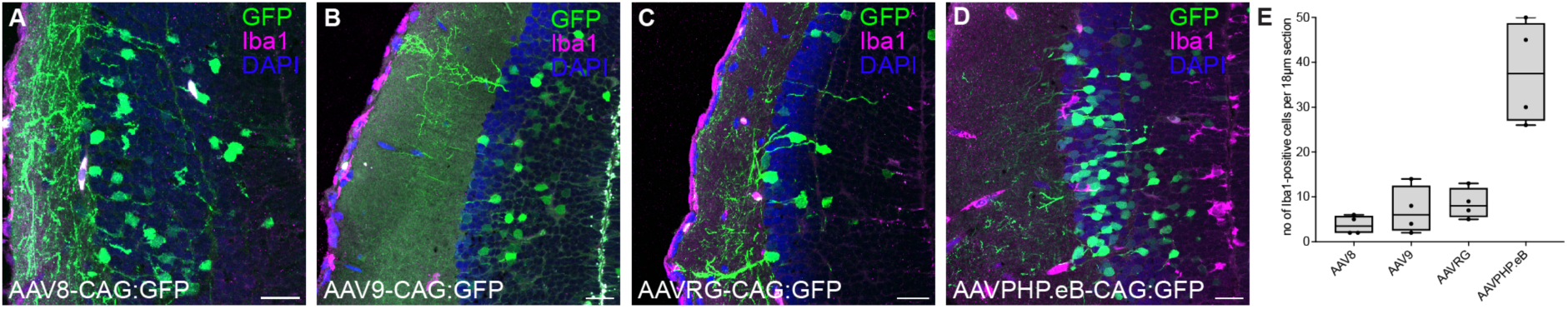
Degree of AAV-induced immune cell infiltration varies based on serotype. (A-D) Confocal images of cryosections of brains injected with either AAV8-CAG:GFP (A), AAV9-CA:GFP (B), AAVRG-CAG:GFP (C) or AAVPHP.eB-CAG:GFP (D) respectively. Sections were labeled with anti-Iba1 (magenta). The fluorescent signal for GFP (green) was detected without antibody enhancement. Nuclei are counterstained with DAPI (blue). All scale bars are 50 µm. (n = 2 axolotl for each serotype) (E) Quantification of the number of Iba1-positive cells per section. Boxplot indicates “minimum,” first quartile, median, third quartile and “maximum”.

### Retrograde transport efficiency and mapping of the optic tectum-retina axis

AAVs can be transported in different directions in neurons, which makes it possible to determine input and output of brain regions of interest. Different AAV serotypes can undergo both anterograde and retrograde transport, however the natural ability for retrograde transport is rather low^27^. Therefore, retrograde tracing-specific variants, such as AAVRG have been engineered^28^. We sought to determine the transport direction of the different AAV serotypes. We decided to use the optic tectum and its anterograde input from the retina via the optic nerve^2^ as an ideal test circuit. First, we assessed the ability and efficiency of retrograde transport. We injected AAV8-CAG:GFP, AAV9-CAG:GFP and AAVRG-CAG:tdTomato into the right optic tectum hemisphere and harvested the left retina as well as the brain at four weeks post injection (Figure 4A). Using wholemount immunostaining of both tissues we aimed to determine the infection efficiency of the optic tectum as well as retrograde transport into the retina (Figure 4B-E). While we found efficient infection of the optic tectum using all three serotypes (Figure 4C-E) the number of GFP/tdTomato-positive retinal ganglion cells was different for each serotype. We found that AAV8-CAG rarely labeled retinal ganglion cells (Figure 4B,C) but AAV9-CAG (Figure 4B,D) and AAVRG-CAG (Figure 4B,E) showed retrograde spreading properties. The efficiency and consistency of AAVRG-CAG to label retinal ganglion cells was however higher than that of AAV9-CAG. These results show that both AAV9-CAG and AAVRG-CAG can be retrogradely transported in neurons from the brain to the retina and that both serotypes can be used for assessing retrograde projections.

**Figure 4.**
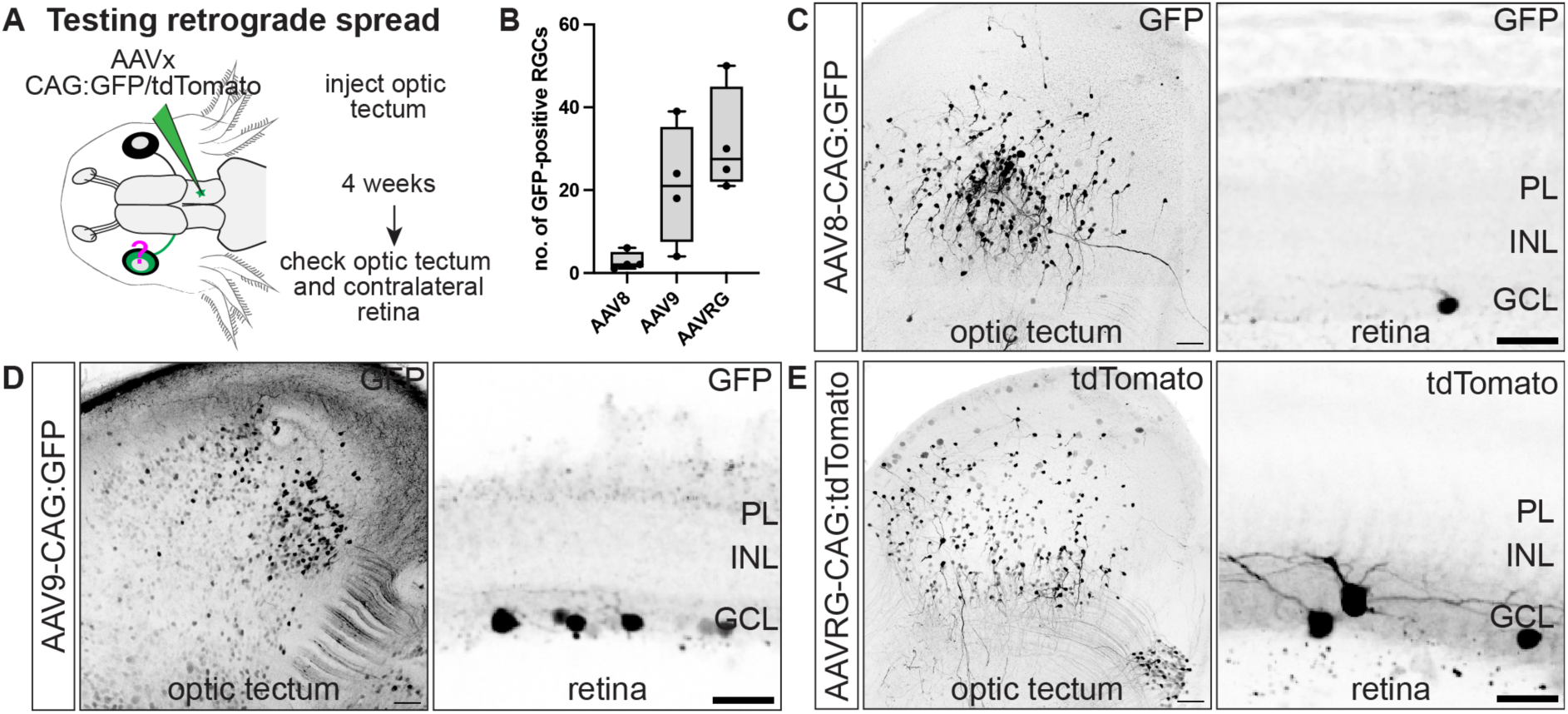
Retrograde spread properties of AAVRG, AAV8 and AAV9 in the axolotl brain. (A) Schematic of experimental paradigm for testing AAV retrograde spread. AAV solution was injected into the right optic tectum hemisphere and the left (contralateral) retina was analyzed for the presence of cell bodies 4 weeks later. (B) Quantification of GFP-positive cell bodies in the retinal ganglion cell layer shows that AAVRG allows for efficient retrograde tracing while AAV8 and AAV9 are less efficient. RGCs-retinal ganglion cells. (n = 4 axolotl for each serotype). Boxplot indicates “minimum,” first quartile, median, third quartile and “maximum”. (C-E) Optical sections from wholemount imaged retina and optic tectum of AAV8-CAG:GFP (B), AAV9-CAG:GFP (C) and AAVRG-CAG:tdTomato (D) respectively. Samples were labeled with anti-GFP (black) or anti-RFP (black). All scale bars are 50 µm. PL-Photoreceptor layer, INL-inner nuclear layer, GCL-ganglion cell layer

### AAV-mediated transduction of the axolotl retina

Next, we wanted to assess the ability of AAV serotypes to infect cells of the retina to ultimately determine anterograde spreading. It has been demonstrated previously that retinal tropism and transduction efficiency vary depending on the serotype and the route of delivery^29^. We decided to inject AAVs intravitreally as this was the most feasible and reproducible method in small axolotl eyes. We injected AAV8-CAG-GFP, AAV9-CAG-GFP and AAVRG-CAG-GFP into the vitreous of the right retina and waited for 4 weeks to reach a high level of expression (Figure 5A). We then fixed both retina and brain and analyzed for GFP expression focusing on which cell types of the retina had been infected and if optic nerves were labeled. We found that AAV8-CAG did not show any ability to transduce retinal cell types (Figure 5B, D). In contrast to that AAV9-CAG showed very efficient labeling of major retinal cell types, including retinal ganglion cells (Figure 5C, D). AAVRG-CAG showed labeling of retinal ganglion cells, however the efficiency was much lower compared to AAV9-CAG (Figure 5C, D). When we analyzed the respective contralateral optic tectum hemisphere for labeling of projections from the optic nerve and found as expected no labeling in AAV8 transduced animals. In AAVRG-CAG injected animals we also did not detect any labeling of the optic nerve. In contrast all AAV9-CAG injected animals showed labeling of the optic nerve (Figure 5F, G). In addition to labeling retinal ganglion cells, we found that AAV9-CAG also labels other retinal cell types, including photoreceptors, amacrine cells, horizontal cells as well as retinal pigmented epithelial cells (Figure 5G). Together, this shows that AAV9-CAG is efficient in transducing retinal cell types and can be used to trace retinal input into the brain.

**Figure 5.**
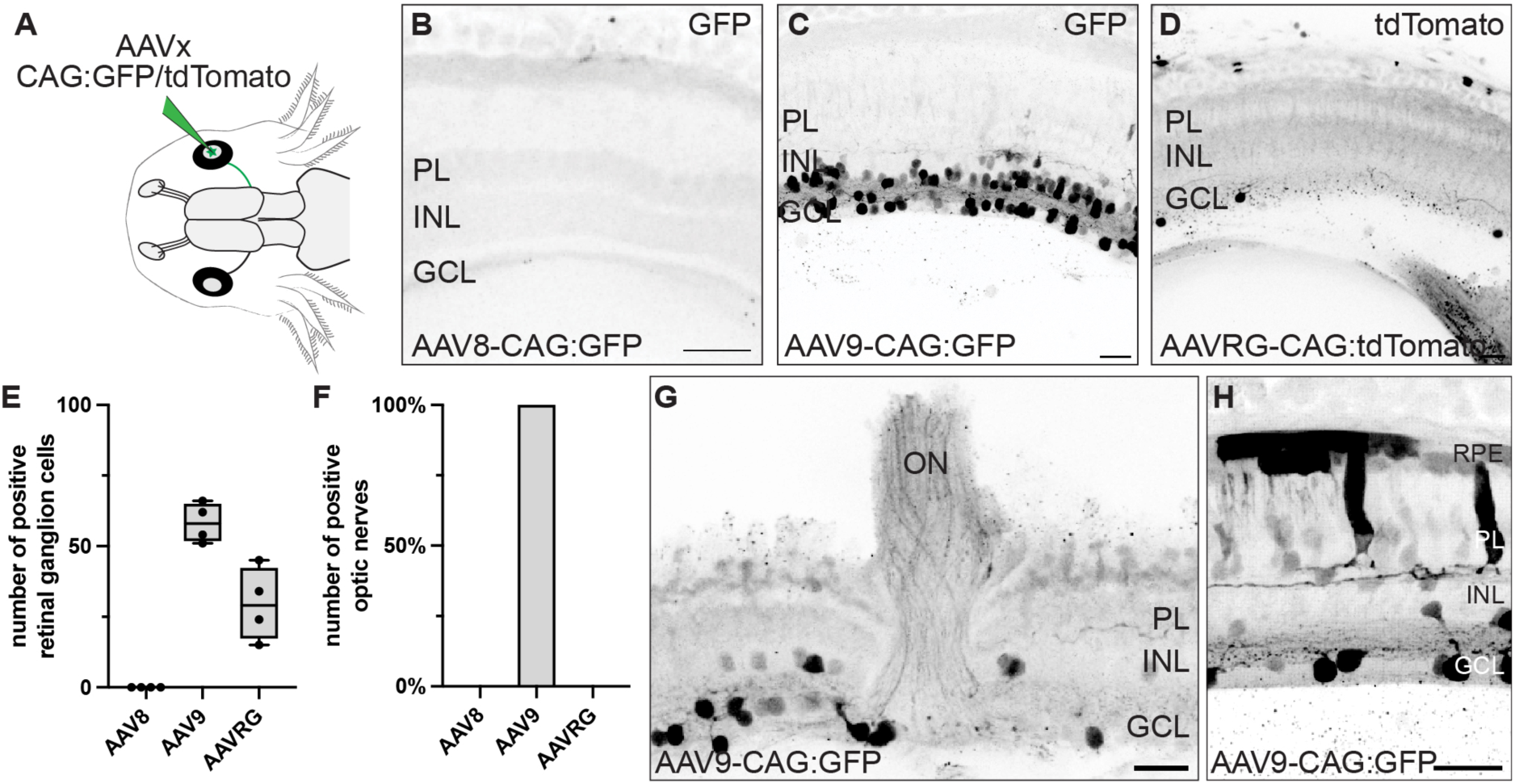
Efficient retinal transduction using AAV9-CAG. (A) Schematic of experimental paradigm for testing AAV anterograde spread. AAV solution was injected into the right retina and the left (contralateral) optic tectum hemisphere was analyzed for the presence of neuronal projections 4 weeks later. (B-D) Optical sections from wholemount imaged retina and optic tectum of AAV8-CAG:GFP (B), AAV9-CAG:GFP (C) and AAVRG-CAG:tdTomato (D) respectively. Samples were labeled with anti-GFP (black) or anti-RFP (black). All scale bars are 50 µm. PL-Photoreceptor layer, INL-inner nuclear layer, GCL-ganglion cell layer, TN-tectal neuropil, ON-optic nerve, RGC-retinal ganglion cell (E) Quantification of GFP-positive retinal ganglion cells shows that only AAV9 and AAV8 allow for efficient ganglion cell labeling. (n = 4 axolotl for each serotype). Boxplot indicates “minimum,” first quartile, median, third quartile and “maximum”.(F) Quantification of GFP-positive cell projections in the optic tectum neuropil shows that sAAV9 allow for efficient anterograde tracing while AAVRG is less efficient and AAV8 does not label any optic nerves. (n = 4 axolotl for each serotype). Boxplot indicates “minimum,” first quartile, median, third quartile and “maximum”. (G-H) Optical sections from wholemount imaged retina transduced with AAV9-CAG:GFP. Sample was labeled with anti-GFP (black). All scale bars are 50 µm. RPE-retinal pigmented epithelium, PL-Photoreceptor layer, INL-inner nuclear layer, GCL-ganglion cell layer, ON-optic nerve

### Tracing of the retina-brain circuitry using AAV9

After determining that AAV9-CAG showed high efficiency to infect retinal ganglion cells and anterograde spread in the optic nerve we decided to trace the retina brain circuitry in the axolotl using this serotype (Figure 6A,B). Optic nerve input into the brain has been determined in other salamander species by use of Golgi staining and horse radish peroxidase mediated tracing^2^ and we thought this to be a suitable comparison to understand how well AAV9-CAG-mediated tracing could resolve those projections. In salamanders, fibers of the optic nerve have been found to distribute to five fields in the brain, the optic tectum, the pretectal nucleus, the area ventrolateralis pedunculi, the thalamus and the hypothalamus^2^. Using AAV9-CAG anterograde tracing we could detect fibers projecting into all these expected brain regions (Figure 6C-G). In addition, we also found GFP-labeled cell bodies in a region of the brain indicating that these neurons were retrogradely labeled (Figure 6H). Using the axolotl brains atlas^30^, we identified this region as the preoptic nucleus. As we had seen before that AAV9-CAG also shows the ability for retrograde spreading this result indicates that these neurons project into the retina. In reptiles and mammals such connections are described as the retinopteal system^31,32^. In other salamander species (*Salamandra salamandra*, *Triturus vulgaris* and *Triturus crestus*) such retinopetal projections have been found^33^. Using AAV9-CAG-mediated tracing we could confirm that retinopetal projections from the brain to the retina are also conserved in the axolotl. Thus, using AAV9-CAG we could faithfully trace all known input from the retina to the brain and additionally uncover the existence of the conserved retinoptal system in the axolotl.

**Figure 6:**
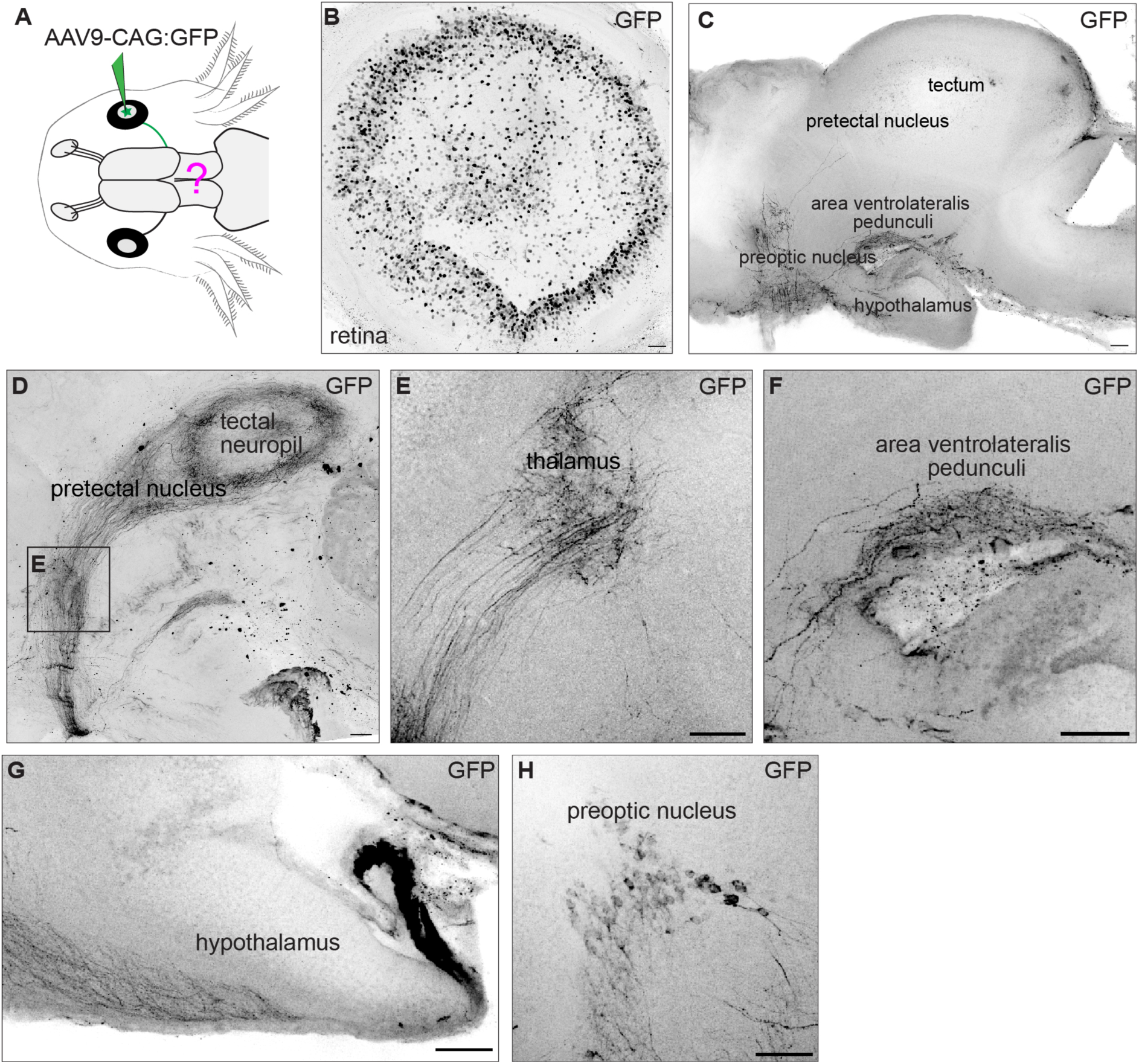
Tracing the retina-brain circuit using AAV9-CAG. (A) Schematic of experimental paradigm. (B) Wholemount maximum intensity projection of retina transduced with AAV9-CAG:GFP. Sample was labeled with anti-GFP (black). Scale bar is 100 µm. (n = 4 axolotl) (C) Wholemount maximum intensity projection of the brain after retinal transduction with AAV9-CAG. Sample was labeled with anti-GFP (black). Scale bar is 100 µm. (D-G) Wholemount maximum intensity projection of brain regions with neuronal projection from retina after retinal transduction with AAV9-CAG. Sample was labeled with anti-GFP (black). All scale bars are 100 µm. (H) Wholemount maximum intensity projection preoptic nucleus region with retrogradely labelled cells. Sample was labeled with anti-GFP (black). Scale bar is 100 µm.

## Discussion

In this study we have investigated the functionality and efficiency of AAVs (AAV1, AAV2, AAV5, AAV8, AAV9, AAVRG and AAVPHP.eB) to transduce cells in the axolotl brain and retina. We found that except AAV2 all tested serotypes transduced cells of the axolotl brain. AAV8, AAV9, and AAVPHP.eB were the most efficient viral vectors resulting in the most labeled cells per area. However, AAVPHP.eB showed an increased immune cell infiltration of the injected brain. Using the CAG promoter to drive expression of fluorescent proteins, we find that neurons rather than ependymoglia are preferentially labeled by AAV8, AAV9, and AAVPHP.eB. Finally, we demonstrate the use of AAV9 to label input and output projections of the retina through anterograde and retrograde tracing. This tracing approach led us to detect a cell population which projects from the brain to the retina.

We determined that in the axolotl AAV9 worked efficiently to transduce both brain and retinal neurons. Especially in the retina, AAV9 showed the highest potential when compared to AAV8 and AAVRG. Similarly, to what has been observed in mouse retina we also found AAV9 to transduce retinal ganglion cells, amacrine cells, horizontal cells, and photoreceptors^34^. AAV9 is of high interest because this serotype is widely used in rodents, especially because of its capacity to cross the blood–brain barrier^35^. Furthermore, AAV9 is one of the few serotypes described to have anterograde transsynaptic spreading ability when used to express Cre recombinase and injected into loxP reporter mice^36^. We have not tested these two abilities of AAV9 in the axolotl yet, however the high efficiency of AAV9 as well as the availability of loxP reporter animals will make this possible in the future.

AAV9 injection in the axolotl retina has allowed us to map the circuitry between retina and the brain and we could recover all known projections of the optic nerve. Additionally, due to the retrograde spreading abilities of AAV9 we discovered the existence of retinopetal cells in the axolotl. Projections from the brain to the retina have been found in other salamander species^33^ and are also known to exist in lampreys^37^, some teleost fish^38^, reptiles^31^, birds^39^ and mammals^32^. While their projection patterns to the inner nuclear layer have been mapped in the retina of other species, future investigation of this cell population in the axolotl will reveal the conservation of such projections.

Here, we have used AAVs to express fluorescent proteins in the axolotl brain and the retina. In other species, AAVs are used for transducing neurons with functional reporter constructs, such as GCaMP-based calcium reporters which enable *in vivo* imaging of neuronal activity^40^. Given the continuous evolution of such reporters and the ongoing advancements tool development in neuroscience, AAVs offer the possibility to stay up-to-date with future sensor developments. Their use in the axolotl is particularly advantageous as they circumvent the need for generating transgenic lines. The introduction of AAV-based neuronal labeling into the axolotl nervous system opens possibilities for future applications, including mapping circuit re-establishment after injury and regeneration. Additionally, the potential for transsynaptic labeling, either through transsynaptic AAVs^36^ or by combining AAVs with rabies virus-based monosynaptic retrograde mapping^41^, holds promise for probing circuit establishment and re-establishment in the axolotl with unprecedented resolution. Notably, rabies virus-mediated transsynaptic tracing has recently been successfully implemented in zebrafish^42^. Furthermore, recent advancements, such as AAV-based screens, for example AAV perturb-seq in the mouse brain^42,43^, could potentially be adapted to the axolotl to perform in vivo screens. Taken together, AAVs as gene transfer tools will allow for various experimental opportunities that were previously unapproachable in the axolotl.

## Acknowledgments

We would like to thank the members of the Tanaka lab for discussions, input, and support. We thank the BioOptics facility IMP/IMBA Core Facilities for outstanding service. We thank the animal care team for excellent axolotl care.

K.L. was supported by a Long-Term Fellowship from the Human Frontier Science Program (LT000605/2018-L) and a Marie Sklodowska-Curie fellowship (101033093), E.M.T. was supported by the Special research programme of the Austrian Science Fund (Project F78) and an Advanced Grant of the European Research Council (RegGeneMems, 742046). For the purpose of Open Access, the authors have applied a CC BY public copyright license to any Author Accepted Manuscript (AAM) version arising from this submission.

## Author Contributions

K.L. and E.M.T. designed the research; K.L. performed the research; K.L. analyzed the data; and K.L. and E.M.T. wrote the paper.

## Supplementary Figures

**Fig. S1:**
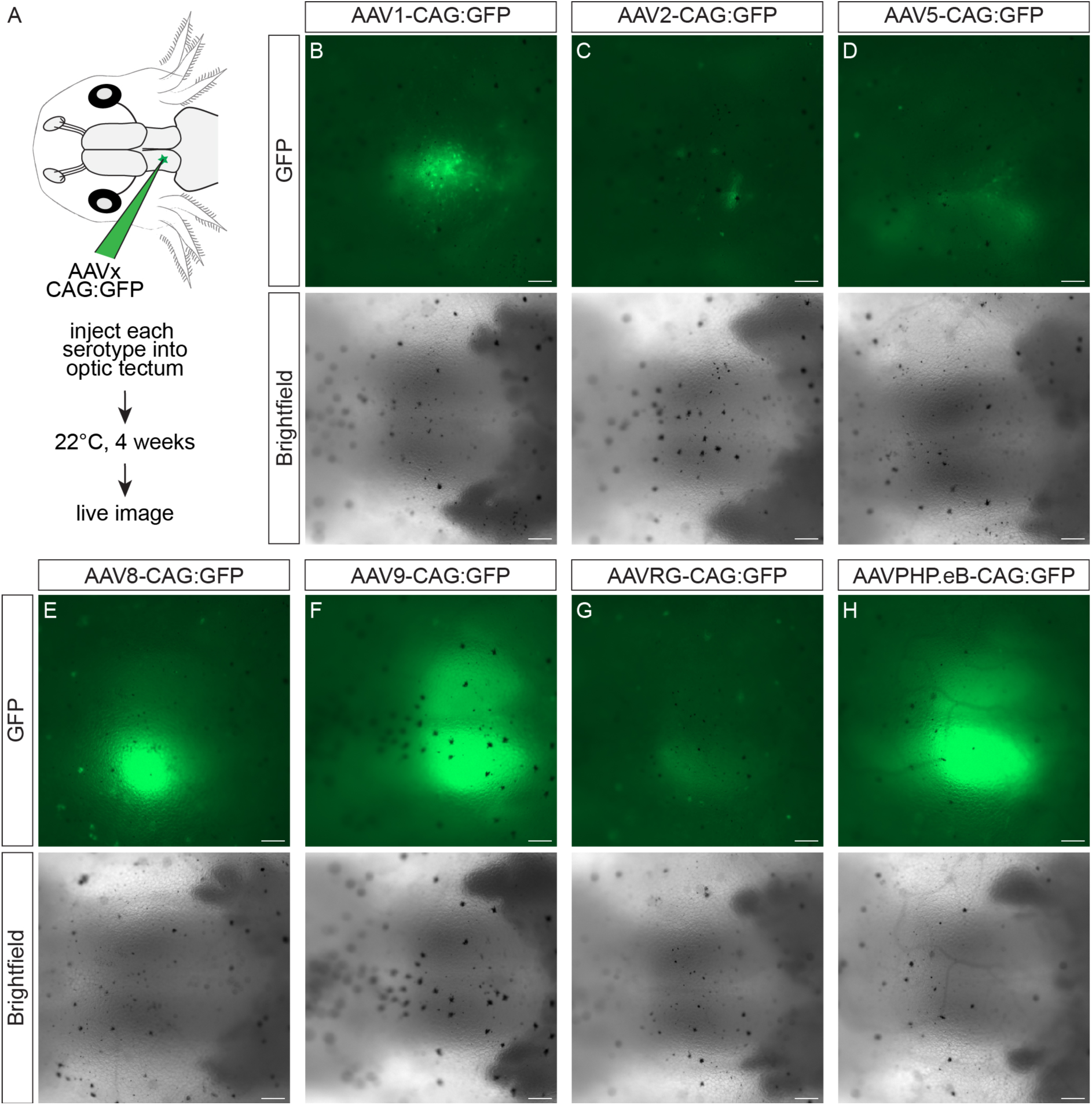
Expression of CAG:GFP using different AAV serotypes in live animals. (A) Schematic of experimental paradigm for AAV serotype efficiency test. Each serotype was injected into the optic tectum neuropil and animals were housed at 22 degrees for 4 weeks until the analysis. (B-H) Live imaging of AAV-CAG:GFP infected brains at 4 weeks after AAV injection. All tested AAV serotypes but AAV2 efficiently transduce axolotl brains and lead to expression of GFP. Upper panels show endogenous GFP fluorescence, lower panels show brightfield images. Scale bars are 250µm. (n = 10 axolotl for each serotype)

**Fig. S2:**
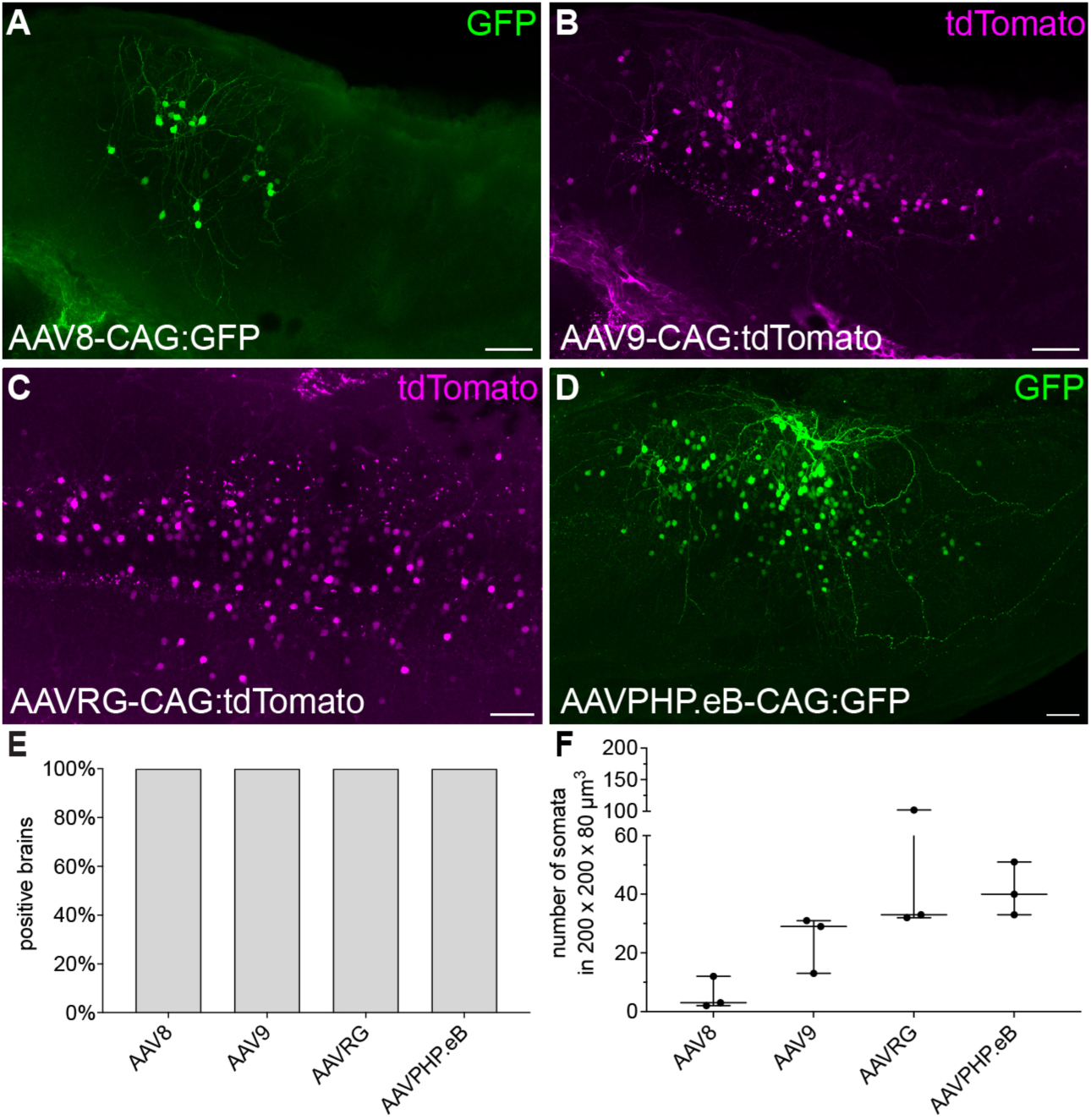
Transduction of cells in juvenile axolotl brains. (A-D) Immunofluorescence for GFP (Green) or tdTomato (magenta) on wholemount, cleared brains of juvenile (10cm nose to tail) axolotls at 4 weeks after AAV8-CAG:GFP (A), AAV9-CAG:tdTomato (B) and AAVRG-CAG:tdTomato (C) and AAVPHP.eB-CAG:GFP (D) respectively injection. Scale bars are 100µm. (n = 3 axolotl for each serotype) (E) Quantification of positive brains 4 weeks after each AAV serotype injection. (F) Quantification of positive somata per 200×200×80 µm^3^.

**Figure S3:**
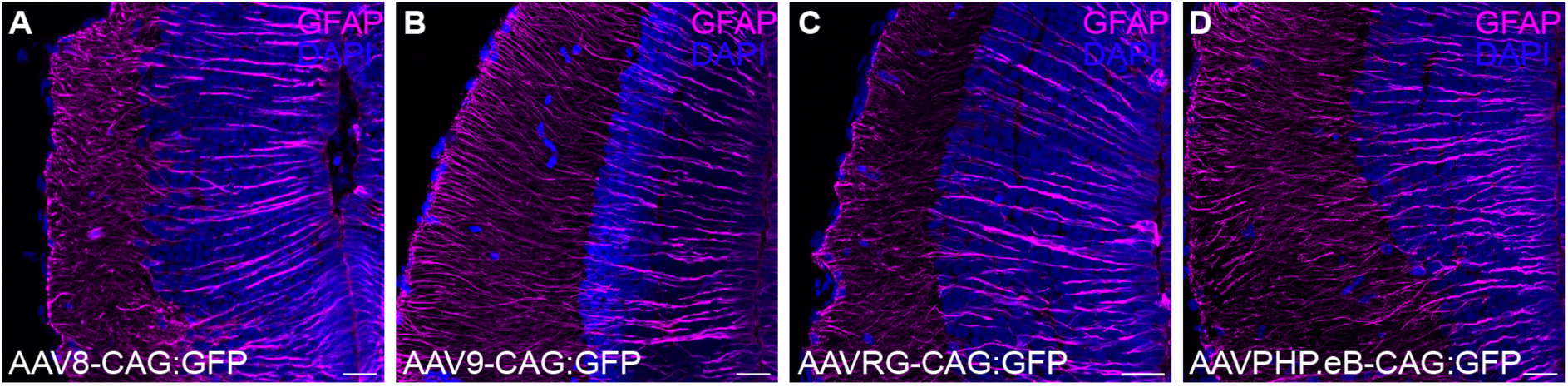
Degree of AAV-induced immune cell infiltration varies based on serotype. (A-D) Confocal images of cryosections of brains injected with either AAV8-CAG:GFP (F), AAV9-CAG:GFP (G), AAVRG-CAG:GFP (H) or AAVPHP.eB-CAG:GFP (I) respectively. Sections were labeled with anti-GFAP (magenta). Nuclei are counterstained with DAPI (blue). All scale bars are 50 µm. (n = 2 axolotl for each serotype)

## Methods

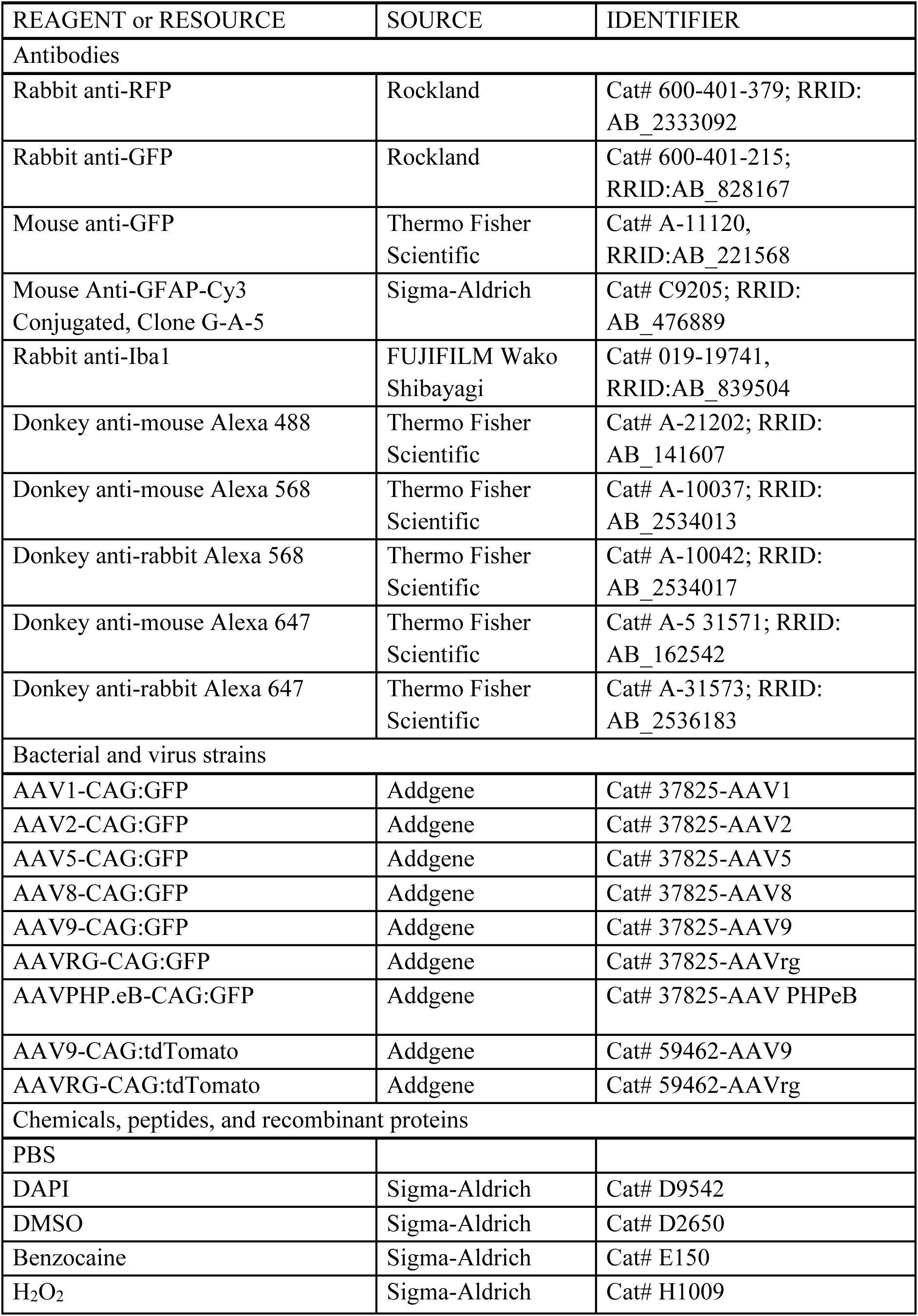
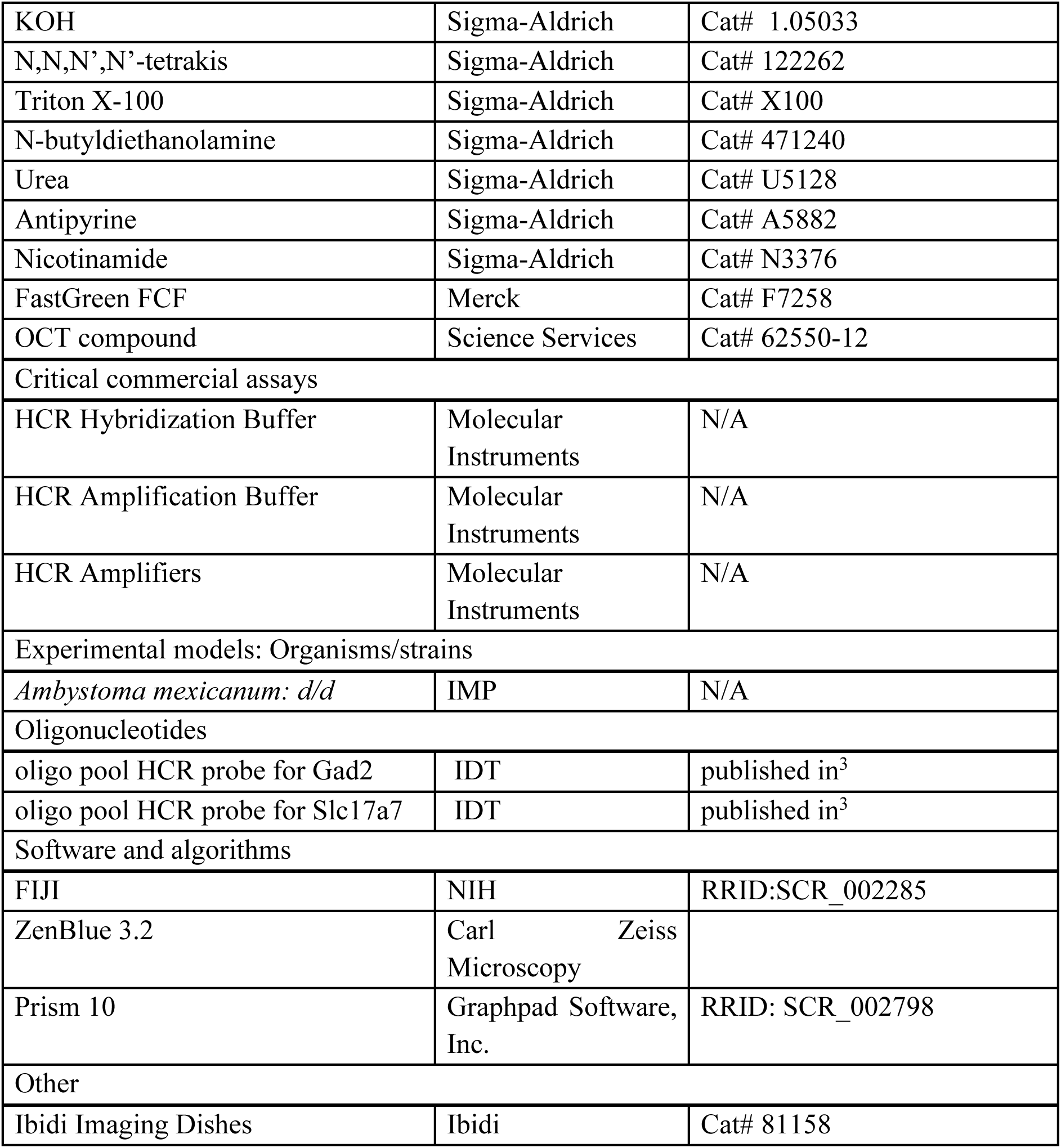
Key Resources Table.

Key Resources Table
| REAGENT or RESOURCE | SOURCE | IDENTIFIER |
| --- | --- | --- |
| **Antibodies** |  |  |
| Rabbit anti-RFP | Rockland | Cat# 600-401-379; RRID: AB_2333092 |
| Rabbit anti-GFP | Rockland | Cat# 600-401-215; RRID:AB_828167 |
| Mouse anti-GFP | Thermo Fisher Scientific | Cat# A-11120, RRID:AB_221568 |
| Mouse Anti-GFAP-Cy3 Conjugated, Clone G-A-5 | Sigma-Aldrich | Cat# C9205; RRID: AB_476889 |
| Rabbit anti-Iba1 | FUJIFILM Wako Shibayagi | Cat# 019-19741, RRID:AB_839504 |
| Donkey anti-mouse Alexa 488 | Thermo Fisher Scientific | Cat# A-21202; RRID: AB_141607 |
| Donkey anti-mouse Alexa 568 | Thermo Fisher Scientific | Cat# A-10037; RRID: AB_2534013 |
| Donkey anti-rabbit Alexa 568 | Thermo Fisher Scientific | Cat# A-10042; RRID: AB_2534017 |
| Donkey anti-mouse Alexa 647 | Thermo Fisher Scientific | Cat# A-5 31571; RRID: AB_162542 |
| Donkey anti-rabbit Alexa 647 | Thermo Fisher Scientific | Cat# A-31573; RRID: AB_2536183 |
| **Bacterial and virus strains** |  |  |
| AAV1-CAG:GFP | Addgene | Cat# 37825-AAV1 |
| AAV2-CAG:GFP | Addgene | Cat# 37825-AAV2 |
| AAV5-CAG:GFP | Addgene | Cat# 37825-AAV5 |
| AAV8-CAG:GFP | Addgene | Cat# 37825-AAV8 |
| AAV9-CAG:GFP | Addgene | Cat# 37825-AAV9 |
| AAVRG-CAG:GFP | Addgene | Cat# 37825-AAVrg |
| AAVPHP.eB-CAG:GFP | Addgene | Cat# 37825-AAV PHPeB |
| AAV9-CAG:tdTomato | Addgene | Cat# 59462-AAV9 |
| AAVRG-CAG:tdTomato | Addgene | Cat# 59462-AAVrg |
| **Chemicals, peptides, and recombinant proteins** |  |  |
| PBS |  |  |
| DAPI | Sigma-Aldrich | Cat# D9542 |
| DMSO | Sigma-Aldrich | Cat# D2650 |
| Benzocaine | Sigma-Aldrich | Cat# E150 |
| H<sub>2</sub>O<sub>2</sub> | Sigma-Aldrich | Cat# H1009 |
| KOH | Sigma-Aldrich | Cat# 1.05033 |
| N,N,N',N'-tetrakis | Sigma-Aldrich | Cat# 122262 |
| Triton X-100 | Sigma-Aldrich | Cat# X100 |
| N-butyldiethanolamine | Sigma-Aldrich | Cat# 471240 |
| Urea | Sigma-Aldrich | Cat# U5128 |
| Antipyrine | Sigma-Aldrich | Cat# A5882 |
| Nicotinamide | Sigma-Aldrich | Cat# N3376 |
| FastGreen FCF | Merck | Cat# F7258 |
| OCT compound | Science Services | Cat# 62550-12 |
| Critical commercial assays |  |  |
| HCR Hybridization Buffer | Molecular Instruments | N/A |
| HCR Amplification Buffer | Molecular Instruments | N/A |
| HCR Amplifiers | Molecular Instruments | N/A |
| Experimental models: Organisms/strains |  |  |
| *Ambystoma mexicanum*: d/d | IMP | N/A |
| Oligonucleotides |  |  |
| oligo pool HCR probe for Gad2 | IDT | published in<sup>3</sup> |
| oligo pool HCR probe for Slc17a7 | IDT | published in<sup>3</sup> |
| Software and algorithms |  |  |
| FIJI | NIH | RRID:SCR_002285 |
| ZenBlue 3.2 | Carl Zeiss Microscopy |  |
| Prism 10 | Graphpad Software, Inc. | RRID: SCR_002798 |
| Other |  |  |
| Ibidi Imaging Dishes | Ibidi | Cat# 81158 |

Key Resources Table

## EXPERIMENTAL MODEL AND SUBJECT DETAILS

### Axolotl

White (d/d) axolotls were used for all experiments. All animals were bred and maintained in IMP facilities and each animal is kept individually. All handling and surgical procedures were carried out in accordance with the local ethics committee guidelines. Animal experiments were performed as approved by the Magistrate of Vienna (Genetically 10 Modified Organism Office and MA58, City of Vienna, Austria, license GZ51072/2019/16 and license GZ665226/2019/21). Axolotl husbandry was performed as described previously (44). Animals of a size of 3cm or 10cm nose to tail were used for all experiments. We are blinded to the animal sex as animals were not genotyped and sexual characteristics are not present at the stages we explored.

## METHOD DETAILS

### Injection of AAVs into brain and retina

AAVs were purchased from Addgene and injected at the maximum concentration (titer ≥ 7×10¹² vg/mL). FastGreen FCF was added for visualization. Before injection, axolotls were fully anesthetized in 0.03% benzocaine. In small (3 cm long) animals the skin and skull are still soft and can be pierced with 27G needle to generate an opening for the injection needle. In big (10 cm long) animals the skin was removed using scissors and forceps and a small incision into the skull was made using a 27G needle. Micropipettes pulled from borosilicate glass were filled with the virus solution and used for pressure injection into the desired region of the brain. In 3 cm animals roughly 0.1µl virus solution per animal and in 10 cm animals 0.5µl virus solution per animal were injected. After surgery, animals were kept in standard holding tanks and the brains were harvested at 4 weeks after injection.

To inject into the retina, a small incision was made in the cornea next to the lens with a micro knife and the borosilicate glass injection needle was inserted into the vitreous. The approximate volume of the vitreous was calculated from measurements with digital calipers as described by Raymond et al. 1988. The vitreous cavity was approximated as the difference between the volume of the entire eyeball minus the volume of the lens. According to this calculation, 0.1µl of virus solution per animal is injected in 3 cm long animals. After 4 weeks the retinae and brains were harvested.

### Brain and eye collection for HCR or immunohistochemistry

For tissue harvesting axolotls were fully anesthetized, decapitated with scissors and brains and eyes were extracted and fixed overnight in 4% paraformaldehyde at 4°C on a horizontal shaker. Fixed brains and eyes were washed six times for 30 minutes each with 1×PBS on a horizontal shaker at 4°C. For further use brains and eyes were either incubated overnight in 30% sucrose in 1×PBS at 4°C if used for cryosectioning or used immediately for wholemount immunohistochemistry. For long term storage, brains and eyes were dehydrated in a methanol/1×PBS series (25%, 50%, 75%, 2 times 100% for 30 minutes each at at 4°C on a horizontal shaker) and stored in 100% Methanol at –20°C.

### Cryosections

Cryoprotected brains (in 30% sucrose in 1×PBS at 4°C) were embedded in Tissue Tek O.C.T. Compound, frozen in liquid nitrogen and stored at –80°C until cryosectioning. Coronal cryosections of 18 μm thickness were prepared from frozen blocks and stored at –20°C until use.

### HCR on cryosections

HCR was performed according to the manufacturer’s instructions by Molecular Instruments with the following modifications: probe concentration was increased to 0.8 pmol and slides were covered with parafilm for all incubation steps. At the end of the procedure the slides were mounted in a glycerol mounting medium (80% glycerol, 1×PBS, 20mM Tris pH 8, 2.5 mg/mL propyl gallate).

### HCR probe design

HCR probe pairs were designed against unique mRNA sequences of the candidate gene. BLAST alignment of the gene sequence against the axolotl transcriptome Amex.T_v47 (50) was used to identify unique sequences. Hybridization wash buffer and amplification buffer, as well as HCR hairpins were purchased from Molecular Instruments. All other probes were ordered as IDT Oligo pools (50pmol). Alexa-546 and Alexa-647 conjugated hairpins were used in this study. Detailed probe sequences have been published previously (ref).

### Immunohistochemistry on cryosections

Cryosections were rehydrated in 1×PBST (0,2% TritonX-100) for 30 min at room temperature. The respective primary antibodies were diluted in 1% normal goat serum (NGS) overnight at 4°C. After six washes for 30 min each at room temperature the secondary antibodies were applied in 1% NGS together with DAPI (1:500 dilution in 1×PBST of 5 mg/ml stock) for 3 hours at 37°C. Slides were mounted with 60% glycerol and stored at 4 °C until imaging.

### CUBIC clearing and wholemount immunostaining of AAV-injected brains and retinae

Clearing and staining of AAV-injected brains and retinae was performed using CUBIC-based clearing using CUBIC-L, CUBIC-R1a and CUBIC-R+(N) solutions (ref) and http://www.cubic.riken.jp). Fixed and washed brains and eyes were bleached in 5%H2O2, 3% KOH in PBST until pigmentation was removed. For delipidation, brains and retinae were incubated in 50% CUBIC-L/R1a solution (mixed 1:1 and diluted to 50% in dH_2_O) for 3 hours on a shaker at room temperature. Afterwards brains and retinae were incubated with 100% CUBIC-L/R1a solution overnight on a shaker at 37°C, followed by washes with 1×PBS for six times 10 min each at room temperature. Respective primary antibodies were used at a dilution as indicated above in 4% Goat Serum, 1% DMSO in 1×PBST and incubated over three nights at 4°C on a horizontal shaker. Secondary antibodies were diluted in 4% Goat Serum, 1% DMSO in 1×PBST, incubated over three nights at 4°C on a horizontal shaker. Brains were washed 6 times 30 min with 1×PBST at room temperature on a horizontal shaker and finally incubated in CUBIC refractive index matching solution CUBICR+(N), on a horizontal shaker until cleared.

### Microscopy

Cleared brains were mounted in Ibidi glass bottom dishes and imaged on an inverted Zeiss LSM980 Axio Observer (inverted) confocal microscope with a 10×/0.3 EC plan-neofluar objective. Cryosections were imaged on an inverted Zeiss LSM980 Axio Observer (inverted) confocal microscope using a 20×/0.8 plan-apochromat objective. ZenBlue 3.2 was used for image acquisition and automatic stitching. Image preparation was performed using FIJI (based on ImageJ1.53c).

## QUANTIFICATION AND STATISTICAL ANALYSIS

### Somata counting

Cell somata counts were performed using the FIJI cell counter plugin. Regions and sections for analysis were selected in a way that they represent the same anatomical location and we performed soma counts in standardized volumes of 200 × 200 × 80 μm^3^ (in 3cm animals) and 200 × 200 × 40 μm^3^ (in 10cm animals) to accommodate for slight variations.

### Statistical analysis

Unpaired t tests were carried out using Prism version 10 (GraphPad; San Diego, CA, USA); p < 0.05 was considered statistically significant.

